# Inference and analysis of population-specific fine-scale recombination maps across 26 diverse human populations

**DOI:** 10.1101/532168

**Authors:** Jeffrey P. Spence, Yun S. Song

## Abstract

Fine-scale rates of meiotic recombination vary by several orders of magnitude across the genome, and are known to differ between species and even between populations. Studying the differences in recombination maps across populations has been stymied by the confounding effect of differences in demographic history. To address this problem, we developed a method that infers fine-scale recombination rates while taking demography into account and applied our method to infer population-specific recombination maps for each of 26 diverse human populations. These maps recapitulate many aspects of the history of these populations including signatures of the trans-Atlantic slave trade and the Iberian colonization of the Americas. We also investigated modulators of the local recombination rate, finding an unexpected role for Polycomb-group proteins and the tri-methylation of H3K27 in elevating recombination rates. Further differences in the recombination landscape across the genome and between populations are driven by variation in the gene that encodes the DNA-binding protein PRDM9, and we quantify the weak effect of meiotic drive acting to remove its binding sites.

---

Meiotic recombination is a fundamental genetic process and a critical evolutionary force which generates haplotypic diversity in sexually reproducing species. In many species, including humans, a zinc finger-containing protein, PRDM9, directs recombination, resulting in hotspots of recombination at its binding sites (*1*). Yet, PRDM9 binds ubiquitously throughout the genome, including at promoters, and only a subset of these correspond to recombination hotspots, suggesting that PRDM9 binding may be necessary but not sufficient (*2*). PRDM9 is capable of trimethylating H3K4 and H3K36 (*3*), and in species that lack a functional copy of *PRDM9*, recombination is concentrated at promoters (*4*), indicating that chromatin structure plays a role in recombination (*5*).

PRDM9-directed recombination has fundamental consequences: recombination hotspots act as breakpoints for chromosomal inheritance, shaping patterns of linked selection (*6*), and an excess of sites where PRDM9 binds one chromosome but not its homolog can lead to male sterility (*7,8*). Such asymmetric binding sites are common in inter-species hybrids, providing a mechanism for the long-known phenomenon of *PRDM9* acting as a speciation gene (*9*). Furthermore, asymmetric binding followed by the introduction of a double-strand break and subsequent homology-directed repair results in meiotic drive against the PRDM9-binding allele, which is equivalent to genic selection at the population level (*10*). Over evolutionary timescales, this meiotic drive erodes the binding sites of PRDM9, generating strong positive selection on *PRDM9* mutants with new binding sites (*11,12*), explaining why *PRDM9* is one of the fastest evolving genes (*13*). These evolutionary dynamics have been studied theoretically (*10,14*) and between species (*12*), but previous empirical investigations have been primarily qualitative rather than quantitative.

We developed a new method, called `pyrho`, to infer fine-scale recombination rates while taking population demography into account and applied it to 26 diverse human populations from phase 3 of the 1000 Genomes Project (1KG) (*15*). We then used the resulting accurate, high-resolution maps to investigate the determinants, impacts, and dynamics of recombination rate variation. Software implementing our method and the inferred recombination maps are available at https://github.com/popgenmethods/pyrho.

Our method uses polymorphism data from unrelated individuals to infer fine-scale recombination maps and can be applied to either phased or unphased data. We make use of a composite likelihood approach (*16–18*) that has been shown to have favorable statistical properties (*19*), but unlike previous methods we avoid computationally expensive Markov chain Monte Carlo (MCMC) by using a penalized likelihood framework and gradient-based optimization. Our approach is between 10 and 450 times faster than `LDhat` (*17*), a popular MCMC-based method, while improving accuracy (Section S1 of Supplementary Material, Figure 1A, Figure S2, and Table S1). We also make use of our recent work on computing two-locus likelihoods (*20*): this allows us to scale to hundreds of individuals whereas `LDhat` can accommodate at most 100 diploid individuals, and, importantly, enables us to account for non-equilibrium demographic histories. Failing to account for past fluctuations in population size has been shown to significantly impact the accuracy of inferred fine-scale recombination rates (*20–22*). The details of our method are presented in Section S1 of Supplementary Material.

**Figure 1.**
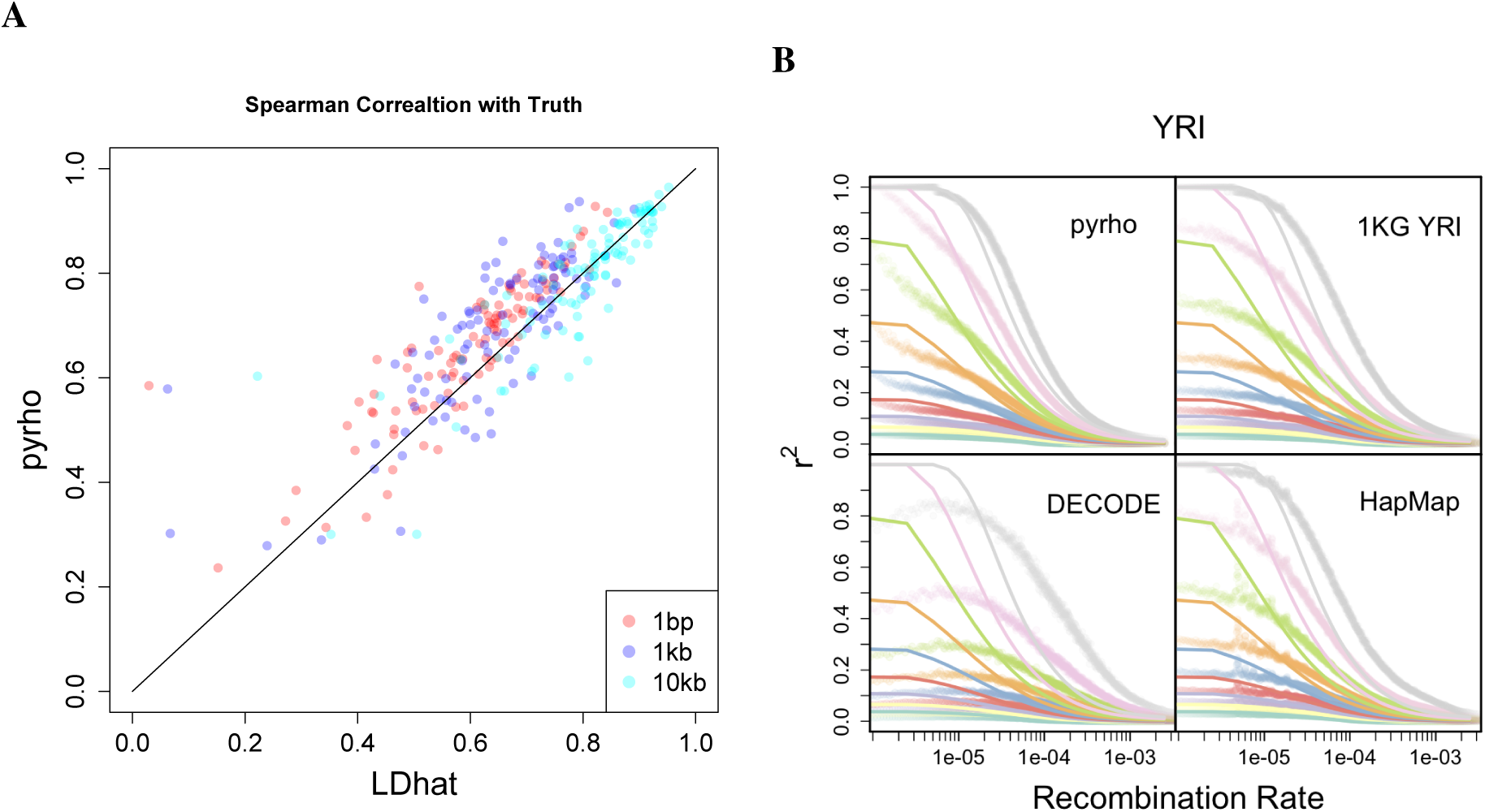
Accuracy of inference on simulated and real data. (**A**) Spearman correlation between inferred and true maps for 100 simulations, each 1 Mb long, for both `pyrho` and `LDhat` with our method showing improved performance especially at finer scales. (**B**) Our inferred recombination maps provide a better fit to observed *r*^2^ values. Solid lines show theoretical deciles of the distribution of *r*^2^ for pairs of sites with minor allele frequency > 0.1 at both sites separated by different recombination distances. Shaded points are the empirical deciles for pairs of sites with minor allele frequency > 0.1 binned by the recombination rate separating them according to different recombination maps.

Using samples of unrelated individuals, we are able to produce more accurate, higher resolution maps from tens to hundreds of individuals than admixture-based (*23,24*) or trio-based methods (*25*), which require data from thousands or tens of thousands of individuals, making our method applicable to a broader set of populations, including unadmixed populations and populations with few sequenced individuals. To explore variation in fine-scale recombination rates across human populations, we inferred population size histories for each of the 26 populations in 1KG (*15*) using `smc++` (*26*) (Figure 2A) and used these size histories to infer population-specific fine-scale recombination maps. Our maps provide a significantly better fit of the observed *r*^2^, a commonly used measure of linkage disequilibrium, especially at finer scales (mean square error between empirical and theoretical quantiles: *p* < 1 × 10^−5^ for each population considered–CEU, CHB, and YRI–for all comparisons between our maps and those inferred in (*15,23,25,27*); twosided permutation test; Section S1.2.3 of Supplementary Material, and Figures 1B and S3). This improvement is particularly pronounced in non-European populations, such as Yoruba (YRI), and could be due to unrealistic assumptions of equilibrium demography made by other methods, a mismatch between the populations used to compute the other maps (e.g., the recombination maps from DECODE (*25*) are inferred using Icelanders), or to previous methods having hyperparameters tuned to European-like demographies.

**Figure 2.**
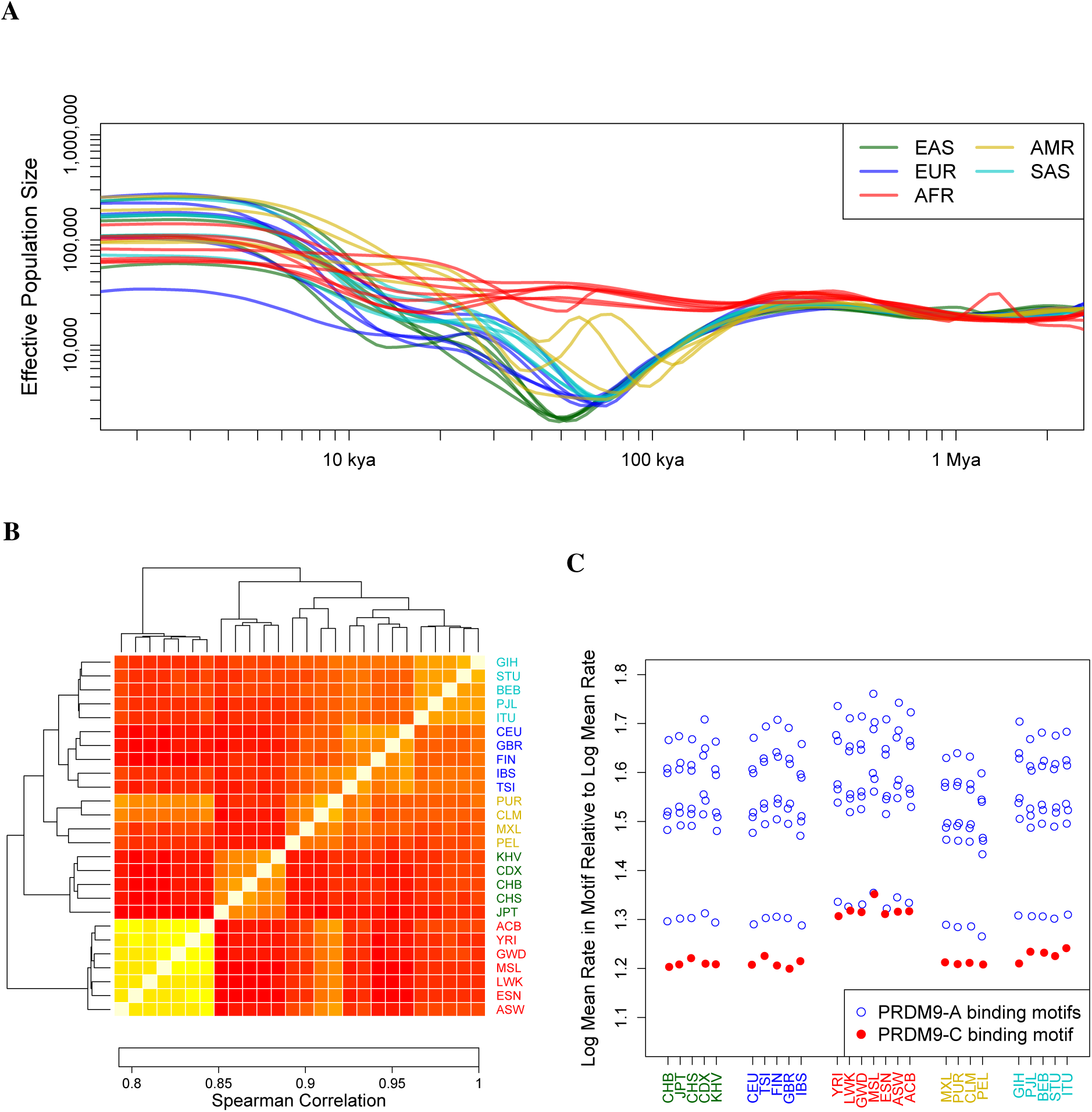
Interplay of demographic history and fine-scale recombination rates. (**A**) Population sizes as inferred by `smc++`. All non-African populations show an out-of-Africa bottleneck, which is deepest in east Asian populations. (**B**) Heatmap of the Spearman correlation between the inferred recombination maps. All maps show a high degree of correlation, yet the relative correlations agree with continental levels of population differentiation. (**C**) Recombination rates at different PRDM9 binding motifs in each population, normalized by the average recombination rate in that population. All PRDM9-A binding motifs show elevated recombination rates across all populations, while PRDM9-C binding motifs have elevated rates in African populations.

Our inferred recombination maps are largely concordant between populations, with high correlation between all maps, even at the single base pair resolution (Spearman’s *ρ* > 0.70 for all pairs), but some differences remain. As seen in Figure 2B, the correlation between recombination maps largely recapitulates known demographic history, clustering continental-level super populations, and at a finer resolution separating northern and southern European populations, and separating the eastern African Luhya (LWK) from west African and primarily west African-descended populations. Admixed American populations show similarity to both African and European populations, particularly the Spanish (IBS), especially in Puerto Ricans (PUR), providing evidence that the trans-Atlantic slave trade and European colonization, respectively, may have impacted the recombination rates of present-day admixed American populations.

While such correlations in fine-scale recombination rates could be due to increased sharing of recombinations in the genealogy of individuals from more closely related populations, they could reflect population-level differences in the determinants of fine-scale recombination rate, such as differences in local chromatin structure, PRDM9 binding site locations, or *PRDM9* alleles. Indeed, there are multiple *PRDM9* alleles that bind different motifs in humans (*28*), and while the *PRDM9*-A allele predominates in all non-African populations, both the the *PRDM9*-A and *PRDM9*-C alleles are common in African populations, suggesting that African populations may have additional recombination hotspots. This is borne out in our inferred maps, with PRDM9-A binding motifs showing elevated recombination rates in all populations but the PRDM9-C binding motif only showing elevated rates in African populations (Figure 2C).

An important consequence of PRDM9-driven recombination is meiotic drive against PRDM9 binding alleles, resulting from homology directed repair of double-strand breaks initiated at the binding motif. While this process has been examined using the divergence between humans and closely related species (*12,29*), the magnitude of the effect has not been quantified. As meiotic drive is equivalent to genic selection on evolutionary timescales (*10*), we may summarize its strength in terms of an effective selection coefficient, *s*, acting against PRDM9 binding alleles. This selection must be strong enough to explain the substantial divergence between humans and closely related species at PRDM9 binding sites (*12,29*), but not so strong as to drive population level differences within humans: male hybrids from species of mice with substantial differences in the locations of PRDM9 binding sites are infertile (*7,8*), whereas such incompatibilities obviously do not exist in humans.

To estimate the selection coefficient *s*, we computationally predicted genomic regions bind PRDM9-A across the genome for each haplotype in 1KG and constructed a diallelic sample frequency spectrum (SFS) for each population by treating sequences that can putatively bind PRDM9-A as one allele and sequences that cannot as the alternative allele (Section S1 of Supplementary Material). Because PRDM9 is predicted to bind ubiquitously and not all PRDM9 binding sites are recombination hotspots, we subdivided each SFS by local recombination rate. We then used each SFS to infer *s* while controlling for background selection and misspecification of the demography (Section S1 of Supplementary Material). For low to moderate recombination rates, we inferred selection coefficients close to zero, consistent with these PRDM9 binding sites not being “true” recombination hotspots, while for the highest recombination rates, we inferred weak but non-zero selection against the PRDM9 binding allele (*s* ≈ 5 − 15 × 10^−5^, Figure 3A).

**Figure 3.**
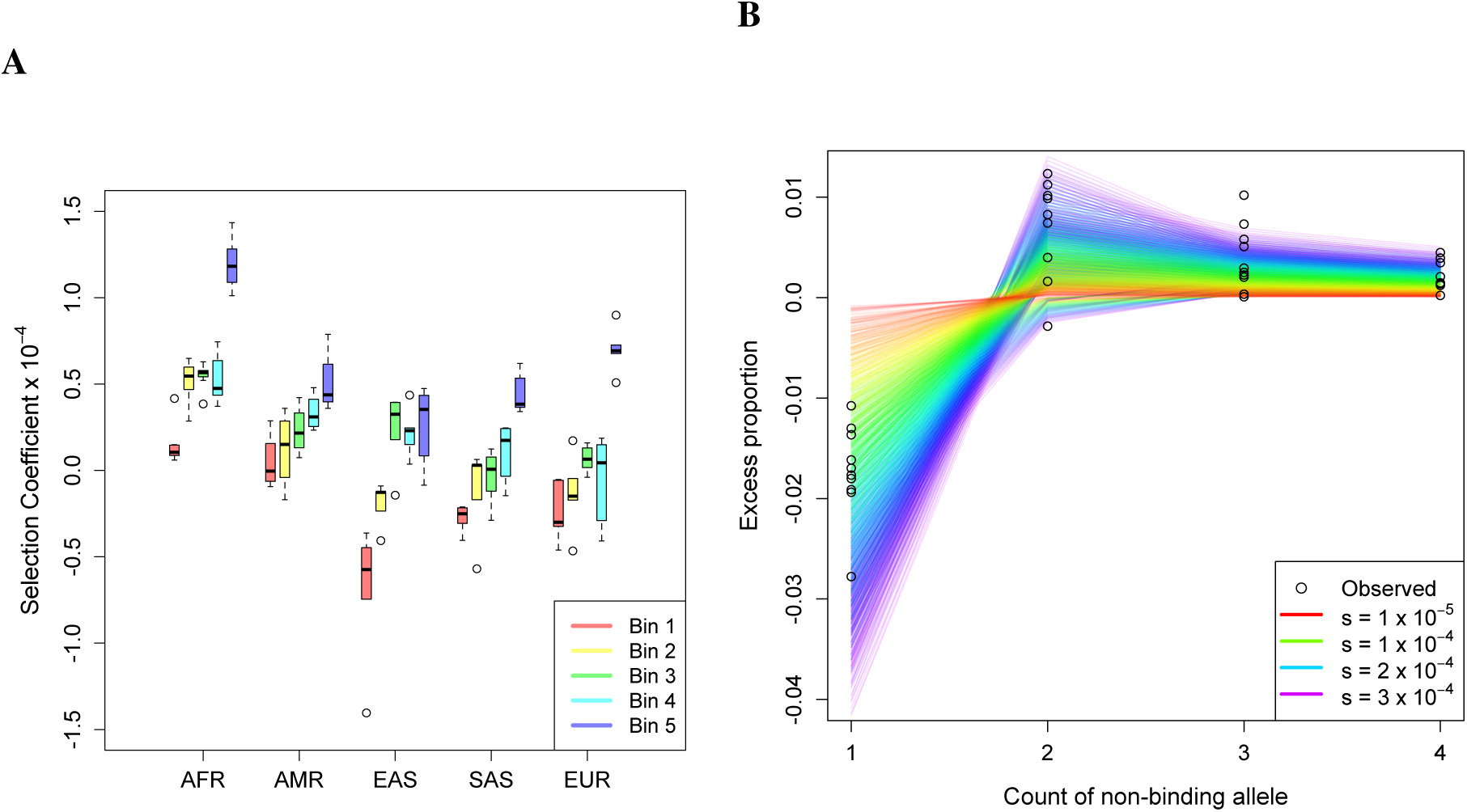
Gene conversion acts like weak selection to remove PRDM9 binding sites. (**A**) Strength of selection acting against PRDM9-A binding alleles for different populations in different bins of recombination rate. Bin 1 is for per-generation rates, *r* ∈ [0, 1.45 × 10^−9^), bin 2 is *r* ∈[1.45 × 10^−9^, 2.78 × 10^−9^), bin 3 is *r* ∈ [2, 78 × 10^−9^, 5.25 × 10^−9^), bin 4 is *r* ∈ [5.25 × 10^−9^, 1.19 × 10^−8^), and bin 5 is *r* ∈ [1.19 × 10^−8^,∞). Bins were chosen such that approximately the same number of polymorphic PRDM9 binding sites fall within each bin. Selection is stronger at bins with higher recombination rates, and is stronger in non-African populations. (**B**) The difference in the frequency of singletons, doubletons, tripletons, and quadrupletons between all PRDM9 non-binding alleles private to European or east Asian populations and SNPs matched by recombination rate. The colored lines are the expectations under a range of demographic scenarios and effective selection coefficients assuming that SNPs evolve neutrally, while the points are the observed differences in the 10 East Asian and European populations. We see a depletion of singletons and excess of more common variants consistent with a selection coefficient between 0.5 × 10^−4^ and 3 × 10^−4^.

The above analysis implicitly assumes that the strength of selection has been temporally constant, which is certainly violated as the motif-determining zinc finger array of *PRDM9* evolves extremely rapidly (e.g., archaic hominins do not have the *PRDM9*-A allele) (*30*). To address this issue, we constructed an SFS of PRDM9 non-binding alleles that are private to Europeans (or private to East Asians) that have most likely arisen since the divergence of Europeans and East Asians (Section S1 of Supplementary Material), and compared the proportion of rare alleles to that seen in a putatively neutral SFS of private SNPs. The observed deficit of singletons and excess of alleles present in more copies (doubletons, tripletons, and quadrupletons) are consistent with an *s* between 0.5 × 10^−4^ and 3 × 10^−4^, suggesting that our previous estimate of *s* is likely a lower bound, but of the correct order of magnitude (Figure 3B).

Because fine-scale recombination rates vary substantially even outside of PRDM9-driven hotspots, we also searched for modulators of fine-scale recombination rates beyond PRDM9, finding a role for chromosome length, distance to the telomere, and chromatin state. Specifically, there is a nearly linear relationship between total physical and total genetic length across chromosomes, with a significantly positive slope and intercept (Figure S4A; slope, *p* = 7.76 × 10^−13^; intercept *p* = 1.30 × 10^−7^). The positive intercept confirms that chromosomes require some minimum number of crossovers during meiosis, while the positive slope indicates that longer chromosomes can and do have more crossovers. Furthermore, recombination rates are elevated in subtelomeric regions (Figure S4B), likely due to the geometry of the chromosomes during meiosis (*31*).

We also found a significant role for chromatin structure in shaping fine-scale recombination rates. We used annotations from chromHMM (*32*) called on 127 ENCODE epigenomes (*33*); because this dataset does not contain calls in gametic cells, we used the most common chromatin state across the 127 cell types as the label for each locus. We found that recombination rate varies significantly across chromatin states (Figure 4; ANOVA *p* < 2.2 × 10^−16^), and that this effect is not driven by differences in background selection (Figure S6 and Section S4 of Supplementary Material). Repetitive regions of the genome have the lowest recombination rates, consistent with a previous finding that a motif present in THE1B repeats is associated with lower recombination rates (*2*), and suggesting that recombination suppression in repetitive regions is a broader phenomenon. We also found lower recombination rates in transcribed regions, providing support for the hypothesis that PRDM9 evolved to direct recombination away from functionally important regions. Furthermore, recombination rates are low in “closed” heterochromatic or quiescent regions perhaps because these regions preclude access to the recombination machinery.

**Figure 4.**
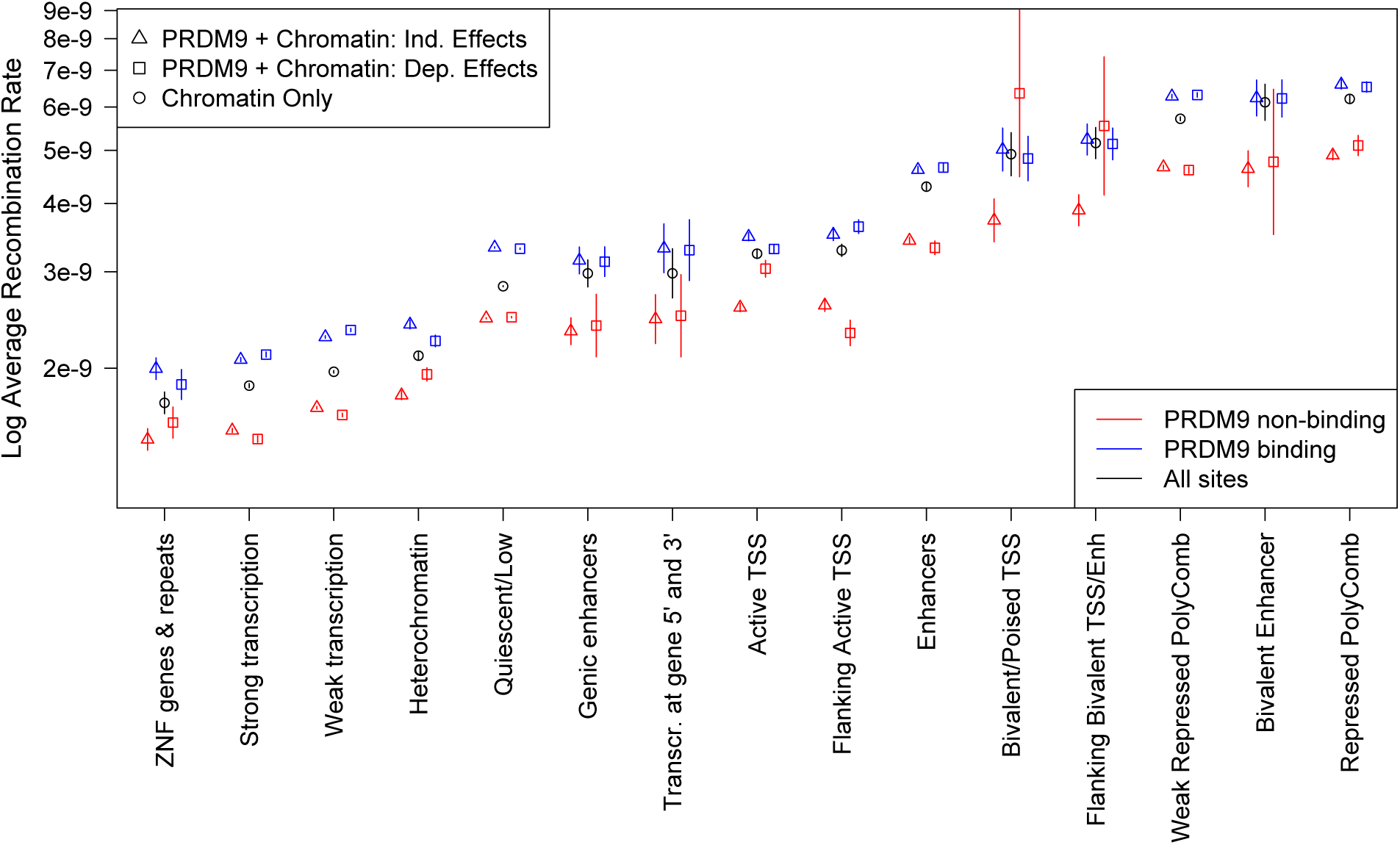
PRDM9 and chromatin structure shape fine-scale recombination rates. Different chromatin states have substantially different average recombination rates as determined by fitting a model using only chromatin state (Chromatin only), a model with independent chromatin state and PRDM9 binding effects (PRDM9 + Chromatin: Ind. Effects), and a model where PRDM9 binding may have a different effect in different chromatin states (PRDM9 + Chromatin: Dep. Effects). Sites characterized by H3K27me3 marks (bivalent states and regions repressed by Polycomb) have the highest recombination rates, while repetitive regions, transcribed regions, and heterochromatic or quiescent regions all have depressed recombination rates.

We found that chromatin states partially characterized by H3K27me3, especially those called as being repressed by Polycomb group proteins (PcGPs), have the highest recombination rates, suggesting a role for H3K27me3 and PcGPs in meiotic recombination. This connection has been noted before, with PcGPs being recruited to double-strand breaks (*34*) and disruption of the PcGP repression pathway leading to improper chromosomal segregation (*35*). This improper segregation in PcGP mutants may be due to a reduced number of successful crossover events in the absence of the H3K27me3 marks deposited by PcGPs. We also note that the substantial impact of chromatin on local recombination rates, along with differences between chromatin structure in male and female gametic progenitor cells, could explain previously observed sex-specific differences in fine-scale recombination rates (*36*). While this manuscript was in preparation, an analysis of a large number of Icelandic trios also found that H3K27me3 and PcGPs are associated with higher local recombination rates (*37*).

The distribution of PRDM9 binding sites across chromatin states is non-uniform (Figure S4D; *χ*^2^ test *p* < 2.2 × 10^−16^) and putative PRDM9 binding is associated with a 49% increase in recombination rate (Figure S4C, *t*-test *p* < 2.2 × 10^−16^), but the variation in recombination rate across chromatin state cannot be explained by differences in PRDM9 binding (*p* < 2.2 × 10^−16^ when controlling for PRDM9 binding status). “Bivalent” chromatin states characterized by active H3K4me1 marks and repressive H3K27me3 marks are particularly enriched for putative PRDM9 binding sites, with over 90% of loci in such states being within 100 bp of a putative PRDM9 binding site. This enrichment cannot be explained by the methyltransferase activity of PRDM9, which trimethylates H3K4 and H3K36 (*3*), leaving the cause of this enrichment unknown.

To investigate the interplay of PRDM9 and chromatin state, we compared a model where PRDM9 affects recombination rate in a chromatin-independent fashion (independent effects model) with a model where PRDM9 can have different effects in different chromatin contexts (dependent effects model), and found that the dependent effects model fits better (*F* -test *p* < 2.2 × 10^−16^). In spite of favoring the dependent effects model, we found that in most chromatin states, the predicted mean recombination rate is similar to that in the independent effects model (Figure 4), indicating that PRDM9 and chromatin state usually act independently. A notable exception is at transcription start sites, where PRDM9 binding is found to have an attenuated effect on recombination rate. This could indicate that the recently discovered ability of PRDM9 to act as a transcription factor may be antagonistic to its role in directing recombination or that PRDM9-independent mechanisms act to suppress recombination at transcriptions start sites (*2*).

## Data and software availability

Software implementation of pyrho and inferred recombination maps can be downloaded from https://github.com/popgenmethods/pyrho.

## Acknowledgments

We would like to thank Ethan Jewett for testing our software and Jane Yu for assistance in downloading and preparing the ENCODE data. This research is supported in part by an NIH grant R01-GM108805, and a Packard Fellowship for Science and Engineering. Y.S.S. is a Chan Zuckerberg Biohub Investigator. This research used resources of the National Energy Research Scientific Computing Center, a DOE Office of Science User Facility supported by the Office of Science of the U.S. Department of Energy under Contract No. DE-AC02-05CH11231.

