## Supplementary Material for "Inference and analysis of population-specific fine-scale recombination maps across 26 diverse human populations"

### S1 Methods and Materials

#### S1.1 Gradient-based estimation of fine-scale recombination rates

##### S1.1.1 Penalized composite-likelihood method for phased data

To infer a fine-scale recombination map using $n$ haplotypes with $L$ SNPs, an obvious first approach would be to attempt to either maximize the likelihood

$$\max_{\rho_1,\cdots,\rho_{L-1}} \mathbb{P}\left[(h_{i\ell})_{(i:1\ldots n),(\ell:1\ldots L)}|\rho_1,\ldots,\rho_{L-1}\right]$$

or obtain a posterior

$$\mathbb{P}\left[\rho_1,\ldots,\rho_{L-1}|(h_{i\ell})_{(i:1\ldots n),(\ell:1\ldots L)}\right] \propto \mathbb{P}\left[(h_{i\ell})_{(i:1\ldots n),(\ell:1\ldots L)}|\rho_1,\ldots,\rho_{L-1}\right]\mathbb{P}\left[\rho_1,\ldots,\rho_{L-1}\right]$$

where $\rho_1,\ldots,\rho_{L-1}$ are the recombination rates between each pair of adjacent SNPs, and $h_{i,\ell}$ is the allele of haplotype $i$ at position $\ell$. Unfortunately, the full-likelihood of the data is intractable. Many methods then make the following approximation (*1*) (but see (*2–4*), which use machine learning or regression approaches to infer recombination rates based on simulations, and (*5, 6*) which use hidden Markov models)

$$\mathbb{P}\left[(h_{i\ell})_{(i:1\ldots n),(\ell:1\ldots L)}|\rho_1,\ldots,\rho_{L-1}\right] \approx \prod_{\ell,k:|\ell-k|<w} \mathbb{P}\left[(h_{i,\ell})_{i:1\ldots n},(h_{i,k})_{i:1\ldots n}|\rho_\ell+\cdots+\rho_{k-1}\right]$$

with some window size $w$, which works well in practice and has attractive theoretical properties (*7*). This also has a justification from the composite-likelihood literature (*8*). Importantly, this pairwise likelihood only depends on the total recombination rate separating the two points, which suggests that one could precompute these likelihoods for each possible two-locus haplotype configuration at a grid of recombination rates. Recent work (*9*) has enabled the computation of these likelihoods for sample sizes in the hundreds.

A drawback of this composite-likelihood approach is that it tends to produce extremely variable estimates. To reduce the variance in the estimate, past approaches have included a prior over recombination maps that explicitly enforces smoothness and have used Markov chain Monte Carlo (MCMC) to obtain samples from the composite posterior over recombination maps (*10–12*). Note that these samples are from a "composite posterior" and not a true posterior, because we have replaced the true likelihood with the composite likelihood. Thus, although these methods provide some sense of the uncertainty in the estimated recombination map, the estimated uncertainty is likely to be inaccurate (*11*). A downside of MCMC is that it is slow due to the need to repeatedly evaluate the composite likelihood.

We circumvent MCMC by performing penalized composite-likelihood inference. To enforce that recombination maps are smooth, but to allow for some large jumps (e.g., at hotspots) we add an $\ell_1$ penalty to the difference of the log of adjacent recombination rates, which is referred to in other settings as the fused-LASSO (*13*). Specifically, we seek to solve the following optimization problem:

$$\max_{\rho_1,\ldots,\rho_{L-1}} \left\{ \sum_{\ell,k:|\ell-k|<w} \log \mathbb{P}\Big[(h_{i,\ell})_{i:1\ldots n}, (h_{i,k})_{i:1\ldots n}\Big|\rho_\ell + \ldots + \rho_{k-1}\Big] - \lambda \sum_{\ell=1}^{L-2} |\log(\rho_{\ell+1}) - \log(\rho_\ell)| \right\}.$$

Note that this is a high-dimensional optimization problem, making derivative-free optimization methods prohibitively slow. We therefore seek to compute gradients of the likelihood with respect to $(\rho_1, \ldots, \rho_{L-1})$, which is problematic because we have replaced exact evaluation of the pairwise log-likelihoods by looking up entries in a precomputed table. To sidestep this issue, we linearly interpolate between the precomputed log-likelihoods, which makes computing gradients an elementary exercise in linear algebra. Note that, due to using linear interpolation, there are non-differentiable points of the likelihood function, but we circumvent this issue by arbitrarily using the slope of the line infinitesimally to the right of any non-differentiable point. We find that this does not affect the results dramatically. Furthermore, for values of the recombination rate that lie outside of the ranges precomputed in the lookup table, we use the closest entry in the lookup table (either the maximum or minimum recombination rate in the table) and treat the derivative as zero. We utilize these gradients in a proximal gradient descent method for fused-LASSO problems (*14*). We found that this optimization scheme usually converges within tens of evaluations of the objective function, making it highly efficient.

One further subtlety is that this optimization problem is non-convex, implying that there may be local optima in which our optimization scheme could get stuck. One could initialize the optimization at a number of random points and then take the best result, but we take an alternate approach. We first perform a univariate minimization, treating the region as having a single, constant recombination rate. We then use this estimate as our initialization, which should further regularize the optima we find toward being "close" to the constant recombination map.

To further speed up inference, we divide the genome into windows that contain 4001 SNPs that overlap by 100 SNPs and optimize each window independently. We then trim the inferred recombination rates corresponding to the first and last 50 SNPs from each window, and combine the resulting estimates to obtain a recombination map. This process of windowing the genome allows us to run many optimizations in parallel.

Our method is implemented in python and makes extensive use of `numba` (*15*), a just-in-time LLVM compiler for python, to optimize numerical routines. We also make use of `cyvcf2` (*16*) to enable the rapid parsing of VCF, bgzipped VCF, and BCF file formats.

#### S1.1.2 Handling unphased data

Our method can also handle unphased data for genotypes from diploid organisms. In principle, one would want to maximize

$$\max_{\rho_1,\cdots,\rho_{L-1}} \sum_{h \text{ consistent with } g} \mathbb{P}\left[(h_{i\ell})_{(i:1\ldots n),(\ell:1\ldots L)}|\rho_1, \ldots, \rho_{L-1}\right],$$

where $g$ is the observed unphased data and "$h$ consistent with $g$" would be the set of phased haplotypes that are equivalent to $g$ when unphased. We could then apply our composite-likelihood approximation to obtain

$$\max_{\rho_1,\cdots,\rho_{L-1}} \sum_{h \text{ consistent with } g} \prod_{\ell,k:|\ell-k|<w} \mathbb{P}\left[(h_{i,\ell})_{i:1\dots n}, (h_{i,k})_{i:1\dots n} | \rho_\ell + \cdots + \rho_{k-1}\right],$$

but unfortunately the outer sum is intractable as it requires phasing all sites simultaneously. Furthermore, it would be difficult to compute gradients under this formulation due to the product. Instead, we make a further approximation by swapping the sum and product to obtain

$$\prod_{\ell,k:|\ell-k|<w} \sum_{h \text{ consistent with } g} \mathbb{P}\left[(h_{i,\ell})_{i:1\dots n}, (h_{i,k})_{i:1\dots n} | \rho_\ell + \cdots + \rho_{k-1}\right].$$

This makes the sum substantially easier because now we only need to phase two loci at a time, and having the product on the outside allows us to take the log and obtain a linear expression

$$\sum_{\ell,k:|\ell-k|<w} \left[\log\left(\sum_{h \text{ consistent with } g} \mathbb{P}\left[(h_{i,\ell})_{i:1\dots n}, (h_{i,k})_{i:1\dots n} | \rho_\ell + \cdots + \rho_{k-1}\right]\right)\right].$$

Furthermore, we may precompute the values inside of the log by using our lookup table of haploid likelihoods at a grid of recombination rates. We then use these new precomputed lookup tables as a drop-in replacement when running our optimization scheme.

### S1.2 Benchmarking

#### S1.2.1 Timing

To obtain timings for our method and `LDhat` for a realistic use-case, we computed the time it took to infer a recombination map for chromosome-scale data. Both methods make use of the same precomputed lookup table of two-locus likelihoods, and so we did not benchmark the creation of those tables, which has been done previously (*9*). Thus, we compare only the amount of time to infer a recombination map. Using `msprime` (*17*), we simulated ten replicates of data matching the length of chromosome 1 with the HapMap recombination map (*18*) and under the demography inferred for CEU, for a sample size of $n = 196$ haploids. Because `LDhat` does not allow for parallelization, we wrote a python script to separate the data into the same overlapping windows used in our method (windows of 4001 SNPs overlapping by 100 SNPs). We ran our method `pyrho` using 32 cores and also used 32 cores to parallelize `LDhat` runs. For the `LDhat` runs, we then used a python script to combine the output of the runs. Because our scripts for splitting and combining the data for `LDhat` are not optimized, we only timed the total runtime of `LDhat` and compared that to the total time `pyrho` required, which is slightly advantageous for `LDhat`. We used the "optimal" hyperparameters for `pyrho` as discussed below and used the default parameters for `LDhat`, which were tuned to a human-like setting. The timings are presented in Figure S1, showing that in our simulations `pyrho` was on average at least 10 times faster than `LDhat`. Yet, when generating the 1KG maps, `LDhat` was run on windows of 2000 SNPs, and the MCMC was run for 22.5 million iterations per window, whereas we used only one million iterations per window in our timing benchmark. Computing the recombination maps for chromosome 1 for 1KG thus likely took between 22.5 and 45 times longer than the results reported here, suggesting that our method is closer to between 225 and 450 times faster.

#### S1.2.2 Accuracy on simulated data

To assess the accuracy of our method, we used `msprime` (*17*) to simulate 100 sequences of 1 Mb with recombination maps randomly drawn from the HapMap recombination map (*18*) under the demography inferred for CEU. We then used the lookup table generated for CEU, which takes demography into account, for pyrho, while using a constant-demography lookup table for LDhat, as is the default for that program. For each simulation, we took the middle 500 kb and computed the correlation between the true recombination map and the inferred recombination map. We computed the Pearson correlation in both natural and log-scale, and also the Spearman correlation. To assess the correlation at different spatial scales, we computed the moving average of the inferred recombination rates with different widths (1 kb and 10 kb). At all scales, we computed the correlation using only every $10,000^{\text{th}}$ rate to avoid issues with auto-correlation.

We also investigated whether the differences between pyrho and LDhat are due to the optimization scheme (i.e., fused-LASSO vs. MCMC) or due to the effect of taking demographic history into account. Using the same simulations as described above, we reran LDhat using the demography-aware lookup table used by pyrho and computed the same measures of correlation between these inferred maps and the true maps. We found that at fine-scales pyrho outperforms LDhat by any measure regardless of whether LDhat used a constant-demography lookup table or the demography-aware lookup table. At broader scales, pyrho outperformed LDhat if LDhat used a constant-demography lookup table, but performed comparably to LDhat using the demography-ware lookup table. Meanwhile, the demography aware version of LDhat outperformed the version of LDhat that assumed a constant-demography at all scales. The results are summarized in Table S1.

#### S1.2.3 Comparison of $r^2$ on the 1000 Genomes Project dataset

To get a sense of accuracy on real data, we computed a measure of linkage disequilibrium, $r^2$, between pairs of nearby SNPs. We used vcftools (*19*) with the --hap-r2, --ld-window 15, --thin 2000, --maf 0.1, and --max-missing 1 flags. Briefly, this removes all missing data, removes SNPs until they are all separated by at least 2 kb, and removes SNPs with a minor allele frequency (MAF) less than 10%, and then computes the $r^2$ for all SNPs within 15 SNPs of each other. For each pair of SNPs we then computed the recombination rate between them as determined by a given fine-scale recombination map. We sorted the pairs of SNPs by the recombination rate between them, and grouped them into bins of 1000 pairs of SNPs, and reported the empirical deciles from that bin. We compared these against the theoretical deciles of the distribution of $r^2$ for sample sizes matched to the observed sample sizes for SNPs with MAF greater than 10%, which we computed from the lookup tables we generated as discussed below. The results for YRI are presented in Figure 1B, and for CEU and CHB in Figure S3.

To determine whether the differences between maps are statistically significant, we computed the mean square error between the empirical and theoretical deciles, averaged across all of the bins of pairs of SNPs for different maps. To compare two maps, we used the difference in their mean square error as a test statistic and obtained a null distribution by performing 1,000,000 permutations of the bins (i.e., randomly assigning each bin to one map or the other, making sure each map has the correct number of bins). We compared the maps that we inferred to the LD-based maps, HapMap (*18*) and 1KG (*20*); a trio-based map, DECODE (*21*); and an admixture-based map (*22*). We performed this comparison for CEU, CHB, and YRI, using the appropriate population for our population-specific maps and the population-specific maps of 1KG. That is, overall we performed 12 comparisons (comparing our maps against 4 others in 3 different populations). For each comparison, we found that our recombination maps had a lower mean square error between the empirical and theoretical deciles with no permutations providing an equal or greater improvement, conservatively implying $p < 1 \times 10^{-5}$ for each comparison.

### S1.3 Inference of population size histories

We applied smc++ (v1.11.1) (*23*) to infer population size histories using a previous build of the genome (hg19). All individuals for a given population were included in the analysis with the first 5 individuals (alphabetically by sample name) being used as "distinguished" individuals in the composite likelihood. We assumed a mutation rate of $1.25 \times 10^{-8}$ per-base per-generation, and masked out sites according to Stephan Schiffels' mappability mask available at https://oc.gnz.mpg.de/owncloud/index.php/s/RNQAkHcNiXZz2fd. Otherwise all default parameter settings of `smc++` were used, and a generation time of 29 years (*24*) was used to convert generations to years.

### S1.4 Lookup table generation

When using the population sizes inferred in the previous section to build lookup tables for `pyrho`, we made some approximations to reduce the computational cost. The population size functions returned by `smc++` using the `plot` command are piecewise constant, with many pieces. To reduce the number of pieces, we started at present and combined adjacent pieces by taking the harmonic mean of the population sizes for those pieces (weighted by their lengths) if all of the pieces that were combined had population sizes within 10% of the resulting harmonic mean. Furthermore, computing the initial stationary distribution of two-locus configurations, which depends on the most ancient population size, is computationally expensive, and so after reducing the number of pieces, the most ancient size was set to 19,067 for all populations. Computing the exact two-locus likelihoods requires $O(n^6)$ time, where $n$ is the sample size, and is too computationally prohibitive for sample sizes in the hundreds for 26 populations. In previous work (*9*), we showed that downsampling approximate two-locus likelihoods for a larger sample size, $N$, results in little loss in accuracy, and these approximate likelihoods may be computed in $O(N^3)$ time and downsampled in $O(N^3 \times (N - n))$ time as well. As such, we used this approximation, with $N = 256$ for each population, downsampling to the observed sample size, which ranged from $n = 122$ to $n = 226$ haploids.

### S1.5 Hyperparameter optimization

Our method has two important hyperparameters, namely the window size $w$, which determines how far apart pairs of SNPs must be before we ignore them, and the $\ell_1$ regularization penalty, $\lambda$, that determines the smoothness of resulting map. Because our method is extremely fast, we were able to optimize these parameters for each population to account for differences in sample size and demography. For each population, we used `msprime` (*17*) to simulate 100 regions, each of 1 Mb in length, with a recombination map randomly drawn from the HapMap recombination map (*18*) and with sample size matching the observed sample size. On this dataset, we then ran our method with all possible combinations of $(w, \lambda) \in \{30, 40, 50, 60, 70, 80, 90\} \times \{15, 20, 25, 30, 35, 40, 45, 50\}$. Our method does not estimate a recombination rate before the first SNP or after the last SNP, so we took the estimated recombination rate in the region between the first and last SNP for each simulation and concatenated them together into a single vector, and did the same with the true recombination maps under which we had simulated. We then computed the Pearson correlation of these vectors in both natural and log scale, and also the Spearman correlation; we also computed these correlations at broader scales by taking our estimates and dividing them into non-overlapping windows of length 10kb or 100kb and concatenating the average recombination rate within each window and doing the same to the true recombination maps. We also computed the squared $\ell_2$ norm between the inferred recombination maps and the true recombination maps in both natural and log scale. We computed all of these quantities for each setting of the hyperparameters. To choose the "best" hyperparameters, we looked at each measure of quality and ranked the hyperparameter settings for that measure (e.g., the hyperparameter setting that produced the smallest square $\ell_2$ norm in natural scale between the estimates and the truth would be ranked 1 for that measure). We then chose the hyperparameter setting that minimized the sum of these ranks over all of the measures we considered. Interestingly, non-African populations tended to have higher values of $\lambda$ and lower values of $w$ than African populations, likely due to the lower SNP density in non-African populations resulting from the out-of-Africa bottleneck.

### S1.6 Prediction of PRDM9-A binding sites and SFS construction

To predict PRDM9-A binding sites, we obtained empirical position weight matrices (PWMs) from (*25*). In (*25*), a number of different motifs are presented, but following that paper we only used their motifs Human1, ..., Human7 as "true" PRDM9-A binding motifs. These PWM matrices describe the probability $p_X(\ell)$ of observing a nucleotide $X \in \{A, C, G, T\}$ for each position $\ell$ in the motif. To determine a cutoff for whether to call a particular sequence as matching a particular binding motif or not, we generated 10,000,000 random nucleotide sequences by sampling each position independently, and drawing $A$ or $T$ with probability $0.3$ and $C$ or $G$ with probability $0.2$, which approximately matches the marginal distribution of nucleotides in the human genome. We then computed the log-likelihood, $\log \mathcal{L}$, of each sequence by

$$\log \mathcal{L}^{(i)} := \sum_{\ell=1}^{M} \log \left[ p_{X_\ell^{(i)}}(\ell) \right],$$

where $X_\ell^{(i)}$ is the nucleotide at position $\ell$ in simulation $i$ and $M$ is the length of the motif. We chose the the 9,999,990$^{\text{th}}$ largest log-likelihood as the cutoff for calling a motif. This is equivalent to an approximate $p$-value of $1 \times 10^{-6}$.

We then called PRDM9-A alleles in each haploid sequence in the 1KG dataset on the hg38 genome build as follows. We considered only diallelic SNPs where all individuals have reported genotypes. Sites with more than two alleles or structural variants were treated as missing. Individuals were treated as having the reference allele at all other positions. Then, starting at the first base in the genome, we computed the log-likelihood, as above, for each motif (or its reverse complement) starting at that position, reporting log-likelihoods that are greater than the empirical cutoff for that motif, and then moving to the next base and repeating. We skipped any starting points where any motif overlapped a missing position. Instead of performing this for each haploid individually, we instead constructed all of the unique haplotypes in the dataset that spanned the region from the starting position to the end of the longest motif, and only computed the log likelihood of each motif on these unique haplotypes.

To construct the PRDM9-A binding site SFS, we took these calls and looked for starting positions where some individuals were called as matching one of the PRDM9-A binding motifs, and other individuals were not predicted to bind any PRDM9-A motif. We then treated binding and non-binding as the two alleles and constructed a standard diallelic SFS. We also constructed SFSs for each population by restricting to only sites with a recombination rate inferred in that population within some range.

### S1.7 Inference of selection coefficients

While a number of software packages exist to fit a selection coefficient to an SFS (e.g., (*26, 27*)), there were a number of peculiarities about the PRDM9 binding SFS that prevented us from using these previous methods; we expect selection to act against PRDM9 binding alleles regardless of whether they are ancestral or derived, and hence we want to "polarize" our SFS by considering the frequency of PRDM9 binding alleles, instead of the frequency of the derived allele or the frequency of the minor allele as is usual. Yet, mutations may act to introduce new PRDM9 binding sites or to disrupt PRDM9 binding, meaning that new mutants may arise at either end of the SFS. To account for this issue, we derived and implemented a method to fit selection coefficients for this particular setting.

Let $\hat{\tau}_n = (\hat{\tau}_{n,1}, \ldots, \hat{\tau}_{n,n-1})$ be the observed PRDM9 binding SFS. That is $\hat{\tau}_{n,k}$ is the number of segregating sites where $k$ individuals have a haplotype that binds PRDM9 and $n-k$ individuals have a haplotype that does not bind PRDM9. As in previous methods (*26, 27*), we fit a selection coefficient by maximizing a multinomial log-likelihood:

$$\log \mathcal{L}_{\text{mult}} \propto \sum_{k=1}^{n-1} \hat{\tau}_{n,k} \log \xi_{n,k}(s, \theta_{\text{bind}}, \theta_{\text{nonbind}}), \tag{1}$$

where $\xi_{n,k}(s, \theta_{\text{bind}}, \theta_{\text{nonbind}})$ is the probability that a segregating site has $k$ binding alleles given a selection coefficient of $s$, a rate $\theta_{\text{bind}}$ of new PRDM9 binding sites appearing via mutation, and a rate $\theta_{\text{nonbind}}$ of all non-segregating PRDM9 binding sites generating a new non-binding PRDM9 allele. As has been shown previously (*28*), we have

$$\begin{aligned} \xi_{n,k}(s, \theta_{\text{bind}}, \theta_{\text{nonbind}}) &= \mathbb{E}_{s,\theta_{\text{bind}},\theta_{\text{nonbind}}}\left[\frac{\hat{\tau}_{n,k}}{\sum_{\ell=1}^{n-1} \hat{\tau}_{n,\ell}}\right] \\ &\approx \frac{\mathbb{E}_{s,\theta_{\text{bind}},\theta_{\text{nonbind}}}\left[\hat{\tau}_{n,k}\right]}{\sum_{\ell=1}^{n-1} \mathbb{E}_{s,\theta_{\text{bind}},\theta_{\text{nonbind}}}\left[\hat{\tau}_{n,\ell}\right]} \\ &= \frac{\mathbb{E}_{s,1,\theta_{\text{nonbind}}/\theta_{\text{bind}}}\left[\hat{\tau}_{n,k}\right]}{\sum_{\ell=1}^{n-1} \mathbb{E}_{s,1,\theta_{\text{nonbind}}/\theta_{\text{bind}}}\left[\hat{\tau}_{n,\ell}\right]}, \end{aligned}$$

where the approximation is exact in the limit of small mutation rates and the final equality follows from the fact that absolute scaling of the mutation rates only determines the total number of segregating sites and not their relative proportions, causing a multiplicative factor to cancel in the numerator and denominator. Therefore, we only need to be able to compute $\mathbb{E}_{s,1,\phi}[\hat{\tau}_{n,k}]$, where $\phi = \theta_{\text{nonbind}}/\theta_{\text{bind}}$. Assuming a panmictic population, this expectation depends on both the unscaled effective population size history, $\eta(t)$, as well as $s$ and $\phi$.

We have thus far suppressed the dependence of this expectation on $\eta$ for notational convenience, but now define $m_{n,k}^{s,\phi}(t)$ to be $\mathbb{E}_{s,1,\phi}[\hat{\tau}_{n,k}]$ for the population size history $\tilde{\eta}(t') = \eta(t + t')$. That is, we truncate the population size history at some point $t$ and treat the resulting function as a new population size history to compute the expectation. Furthermore, define $\mathbf{m}_n^{s,\phi}(t) := (m_{n,1}^{s,\phi}(t), \ldots, m_{n,n-1}^{s,\phi}(t))$. The idea behind our method is to set up and solve a system of differential equations of the form

$$\frac{d}{dt}\mathbf{m}_n^{s,\phi}(t) = g(\mathbf{m}_n^{s,\phi}(t), t)$$

to obtain $\mathbf{m}_n^{s,\phi}(0)$, which is our desired expectation. In the case where $s = 0$, this system of equations turns out to be equivalent to the Moran model (*29*) with a continuous injection of new mutants into classes at the boundary, a result that follows from (*30*) and is further explored in (*31, 32*). That is

$$\frac{d}{dt}\mathbf{m}_n^{0,\phi}(t) = -\big(\mathbf{M}_n(t, 0)\big)^T \cdot \mathbf{m}_n^{0,\phi}(t) - \mathbf{e}_1 - \phi\mathbf{e}_{n-1},$$

where the minus signs arise from our convention of having time run backward, $\mathbf{e}_i$ is the $i^{\text{th}}$ basis vector, and $\mathbf{M}_n(t, s) \in \mathbb{R}^{n-1\times n-1}$ is the well-known generator of the Moran process scaled by the population size:

$$\big(\mathbf{M}_n(t, 0)\big)_{ij} = \begin{cases} -\frac{i(n-i)}{\eta(t)}, & \text{if } j = i, \\ \frac{i(n-i)}{2\eta(t)}, & \text{if } j = i - i, \\ \frac{i(n-i)}{2\eta(t)}, & \text{if } j = i + 1, \\ 0, & \text{otherwise.} \end{cases}$$

In the case where there is selection ($s \neq 0$), there is no closed system of differential equations exactly describing the evolution of this vector (*27, 30*). Yet, it is known that the Moran model with selection converges to the Wright-Fisher diffusion with selection in the limit of large $n$ (*33*). We therefore approximate the dynamics with selection by the Moran process with selection, and we compute these expectations for a larger sample size and then downsample to our observed sample size. With this approximation, we obtain the following system of equations,

$$\frac{d}{dt}\mathbf{m}_n^{s,\phi}(t) \approx -\big(\mathbf{M}_n(t,s)\big)^T \cdot \mathbf{m}_n^{s,\phi}(t) - \mathbf{e}_1 - \phi\mathbf{e}_{n-1},$$

where

$$\big(\mathbf{M}_n(t,s)\big)_{ij} = \begin{cases} -\frac{i(n-i)}{\eta(t)} - s \times \frac{i(n-i)}{n}, & \text{if } j = i, \\ \frac{i(n-i)}{2\eta(t)} + s \times \frac{i(n-i)}{n}, & \text{if } j = i - i, \\ \frac{i(n-i)}{2\eta(t)}, & \text{if } j = i + 1, \\ 0, & \text{otherwise.} \end{cases}$$

Now that we have set up the system of differential equations, we show how to efficiently solve it. We assume that $\eta$ is piecewise constant, with sizes $\eta_1, \ldots, \eta_{T+1}$ and breakpoints $t_1, \ldots, t_T$, setting $t_0 := 0$ for ease of notation. For convenience denote the lengths of the pieces as $\Delta_1, \Delta_2, \ldots, \Delta_T$ where $\Delta_k = t_k - t_{k-1}$ for $k > 1$, and also let $\widetilde{\mathbf{M}}_n(k,s) := \mathbf{M}_n(t,s)$ for any $t$ in the $k^{\text{th}}$ epoch. We begin at the most ancient interval, which runs from $t_T$ to $\infty$ and has size $\eta_{T+1}$. Since this epoch is infinitely long, we can compute $\mathbf{m}_n^{s,\phi}(t_T)$ by finding the stationary distribution of this process. That is, we solve

$$\mathbf{0} = -\big(\widetilde{\mathbf{M}}_n(T+1,s)\big)^T \cdot \mathbf{m}_n^{s,\phi}(t_T) - \mathbf{e}_1 - \phi\mathbf{e}_{n-1}$$

for $\mathbf{m}_n^{s,\phi}(t_T)$, using a sparse linear solver implemented in scipy (*34*).

Now, assume that we have computed $\mathbf{m}_n^{s,\phi}(t_k)$. We may compute $\mathbf{m}_n^{s,\phi}(t_{k-1})$ by separately considering what happens to mass already in the system and what happens to mass that is injected during this epoch. Mass already in the system simply evolves according to $\widetilde{\mathbf{M}}(k,s)$, so the contribution of existing mass is $\exp\left\{\Delta_k\Big(\widetilde{\mathbf{M}}_n(k,s)\Big)^T\right\} \cdot \mathbf{m}_n^{s,\phi}(t_k)$ which can be efficiently computed using `expm_multiply` implemented in scipy (*35*). For newly arising mass, we further condition on when the mass arose, resulting in

$$\int_0^{\Delta_k} \exp\left\{t\Big(\widetilde{\mathbf{M}}_n(k,s)\Big)^T\right\} \cdot (\mathbf{e}_1 + \phi\mathbf{e}_{n-1})dt = \Big(\widetilde{\mathbf{M}}_n(k,s)\Big)^{-T} \exp\left\{\Delta_k\Big(\widetilde{\mathbf{M}}_n(k,s)\Big)^T\right\} \cdot (\mathbf{e}_1 + \phi\mathbf{e}_{n-1}),$$

which can be computed efficiently, again with `expm_multiply` and a sparse linear solver to avoid needing to invert a matrix. Combining, this results in

$$\mathbf{m}_n^{s,\phi}(t_{k-1}) = \exp\left\{\Delta_k\Big(\widetilde{\mathbf{M}}_n(k,s)\Big)^T\right\} \cdot \mathbf{m}_n^{s,\phi}(t_k) + \Big(\widetilde{\mathbf{M}}_n(k,s)\Big)^{-T} \exp\left\{\Delta_k\Big(\widetilde{\mathbf{M}}_n(k,s)\Big)^T\right\} \cdot (\mathbf{e}_1 + \mathbf{e}_{n-1}),$$

and iterating this computation we arrive at $\mathbf{m}_n^{s,\phi}(0)$ as desired.

Finally, to find selection coefficients, we can numerically maximize Equation (1) using Powell's direction set method (*36*) as implemented in scipy.

To minimize the effect of using the Moran process with selection to approximate the Wright-Fisher process, we used a larger sample size of 256, and then downsampled to the desired sample size. We performed this for each population for each window of recombination rates, using the decimated `smc++` inferred population sizes.

We were concerned about potential biases arising from either misspecification of the population sizes or differences in background selection due to differences in the recombination rate. To alleviate this bias, we also computed selection coefficients using only SNPs as a putatively neutral control. We tabulated SFSs for SNPs in the same recombination bins, say $\hat{\tau}_n^{\text{SNP}}$, and then computed

$$\tilde{\tau}_n^{\text{SNP}} = \alpha\hat{\tau}_n^{\text{SNP}} + (1-\alpha)\text{reverse}\left[\hat{\tau}_n^{\text{SNP}}\right]$$

where the reverse$[\cdot]$ operator reverses the indexing of the vector, with $\alpha$ chosen to match the ratio of the 1$^{\text{st}}$ to the $(n-1)^{\text{th}}$ entries of the SFS between the SNP and PRDM9 binding cases. We then inferred selection coefficients for $\tilde{\tau}_n^{\text{SNP}}$ using the same method as described above. Our reported de-biased estimate for a given recombination bin and population are then the selection coefficients inferred for the PRDM9 binding SFS minus the selection coefficient inferred for the matched SNP SFS.

### S1.8 Differences in SFS for private mutations

To compute the expected difference between a normalized SNP SFS and a normalized PRDM9 SFS for mutations that are private to a continental group, we computed the expected SFS as above, but instead of starting with the stationary distribution at the most ancient epoch, we set $\mathbf{m}_n^{s,\phi}(t_{\text{div}})$ to be zero – ignoring all mutations that occurred prior to the divergence time, $t_{\text{div}}$. We then computed the normalized SFS under some selection coefficient $s$ and under neutrality ($s = 0$), and reported the difference. We assumed that $t_{\text{div}}$ was 50 ka ago, but the results are qualitatively similar for all the divergence times we tried between 40 ka and 70 ka ago. To compute the observed difference for Europeans, we looked at only PRDM9 binding alleles (or SNPs) where the binding allele (in the SNP case either SNP allele) was at frequency 1 in all African populations and East Asian populations, pooled across all recombination rate bins. We matched the SNP SFS to the PRDM9 binding SFS in terms of recombination rate as follows. First we partitioned SNPs and PRDM9 binding alleles into different bins based on the recombination rate at the SNP or PRDM9 binding site, and computed an unnormalized SFS for the SNPs within each bin. We then computed an overall SNP SFS by weighting each bin proportional to the number of PRDM9 binding alleles within that bin. The binning scheme we chose was

$$\left(-\infty, \frac{e^{-20}}{40000}\right), \left[\frac{e^{-20}}{40000}, \frac{e^{-19}}{40000}\right), \left[\frac{e^{-19}}{40000}, \frac{e^{-18}}{40000}\right), \ldots \left[\frac{e^{0}}{40000}, \infty\right).$$

We then normalized the PRDM9 binding SFS by this weighted SNP SFS and took the difference. We repeated this procedure swapping the roles of Europeans and East Asians.

### S1.9 Data processing and analysis for determinants of recombination rate variation

For the analyses presented in Figure 4 and Figure S4, we preprocessed the data as follows. We first restricted our analyses to only sites satisfying the previously mentioned mappability mask for which we inferred recombination rates. Then, to partially alleviate issues of spatial dependency, we subsetted these data by taking every 1000$^{\text{th}}$ element. We calculated the expected number of recombinations per chromosome by averaging this subsetted data within each chromosome and then multiplying by the chromosome length. For analyzing the subtelomeres we averaged all entries within the first 10 Mb of each chromosome to obtain an average for the "left subtelomere" and the last 10 Mb for the "right subtelomere". We ignored the missing subtelomeric regions in the acrocentric chromosomes 13, 14, 15, 21, and 22 and only presented results for the right subtelomere for these chromsomes. When analyzing the effect of putative PRDM9 binding or chromatin status, we performed all our analyses in log-space. In our benchmarking, we found that `pyrho` produces errors that are approximately normally distributed in log-space, making the use of $t$-tests, ANOVA, and linear models more appropriate in log-space. All statistical tests were performed in R (*37*).

### S2 Comparison with previous recombination maps

We used LiftOver (*38*) to re-map previously inferred recombination maps to the current genome build (hg38). We compared our maps with maps released with the 1KG project for CEU, CHB, and YRI (*20*); the sex-averaged DECODE recombination map (*21*); the HapMap recombination map (*18*); and the admixture-based maps reported in Hinch *et al.* (*22*) and Wegmann *et al.* (*39*). We then computed correlation (Pearson in natural scale and log-scale, and Spearman, at various spatial resolutions, as described above) between all pairs of maps. The results are presented in Figure S5.

### S3 Effect of genome build

We inferred recombination maps on both the current genome build (hg38) and the previous genome build (hg19) to explore the effect of using LiftOver (*38*) to move recombination maps from one coordinate system to another. This is common in practice, with, for example, the DECODE map being originally called on hg18 (*21*) but commonly used on hg19 following LiftOver. There appears to be only a modest overall effect: even at the single base-pair resolution the Spearman correlation between maps inferred on hg38 and those inferred on hg19 and lifted to hg38 ranged from $\rho = 0.986$ to $\rho = 0.998$ across all populations. Similarly the Pearson correlation in log-space varied from $r = 0.984$ to $r = 0.998$. The Pearson correlation in natural scale was somewhat less reliable, however, ranging from $r = 0.474$ to $r = 0.987$, likely due to the extreme leverage of hotspots in natural scale. The results are summarized in Table S2.

### S4 Effect of background selection on inferred recombination rates

To investigate the effect of background selection on our inferred recombination rates, we downloaded a genome-wide measure of background selection (B-statistics) from (*40*). B-statistics range in value from 0 to 1,000 and reflect the relative loss in genetic diversity as a result of background selection, with 0 being a total loss in diversity and 1,000 representing the truly neutral level of genetic diversity. The available B-statistics are reported in terms of coordinates on hg18, so we used LiftOver (*38*) to re-map the coordinates to hg38. We took the data as processed in Section S1.9 and further restricted to sites with reported B-statistics. We repeated the analyses of the effect of putative PRDM9 binding and chromatin state while controlling for background selection by including the B-statistics as linear covariates. The results are presented in Figure S6. While we observe a high correlation between the inferred recombination rate and the B-statistics (Spearman's $\rho = 0.375$, $p < 2.2 \times 10^{-16}$), the overall impact of chromatin state and PRDM9 binding remains comparable whether or not we control for B-statistics.

Note that B-statistics were originally computed by fitting distributions of selection coefficients for exonic and non-exonic regions to observed patterns of diversity (*40*). Importantly, the impact of these distributions on the diversity at linked neutral sites depends on the genetic distance between the selected site and the neutral site, and hence requires knowledge of the fine-scale recombination map. Due to this circularity, it is difficult to determine whether the observed correlation between B-statistics and our inferred recombination rates is due to lower recombination rates directly causing higher levels of background selection (and hence lower inferred recombination rates being associated with lower B-statistics), or if higher levels of background selection result in a lower apparent effective population size, resulting in underestimated recombination rates. It may be possible to disentangle background selection from changes in local recombination rate by jointly inferring B-statistics and fine-scale recombination rates, but we leave such an undertaking for future work.

### S5 Results on unphased data

To test the performance of our method on unphased data, we performed the hyperparameter optimization described above for each population with genotype data from diploid individuals. We then tested our method on the same benchmarking data used in Section S1.2.2 using the optimal hyperparameters for CEU. The results are presented in Figure S7. For the most part inference using unphased individuals results in performance indistinguishable from that on perfectly phased data. As such we recommend using genotype calls when phasing may be inaccurate.

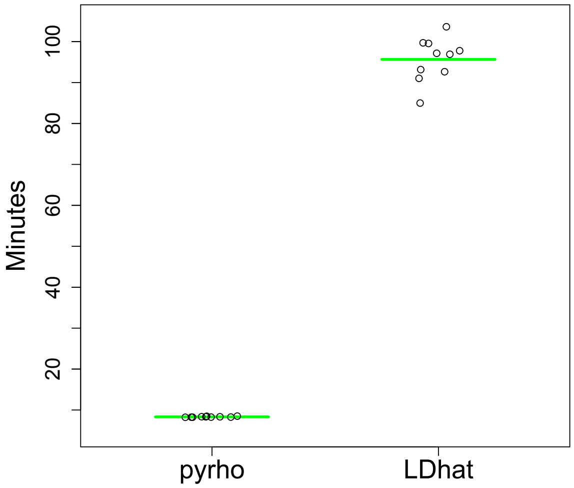


**Figure S1:** Time in minutes to infer a recombination map for a simulated chromosome 1 with 196 haploids, using 32 cores for 10 replicate simulations.

**A** **B**

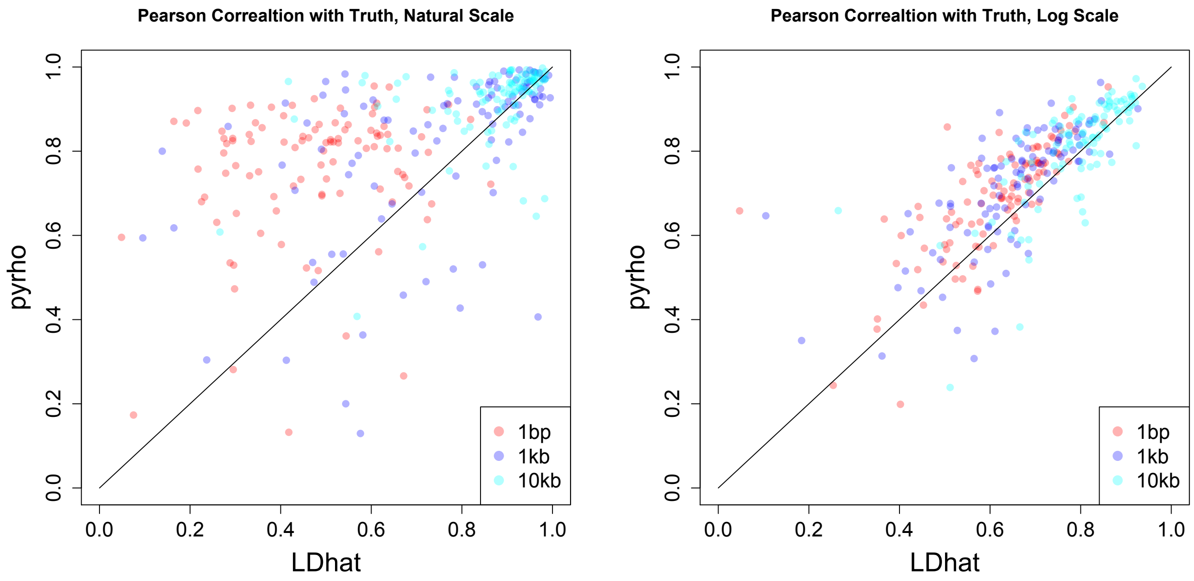


**Figure S2:** Additional measures of accuracy on simulated data. (**A**) Pearson correlation with the truth in natural scale and (**B**) Pearson correlation with the truth in log scale.

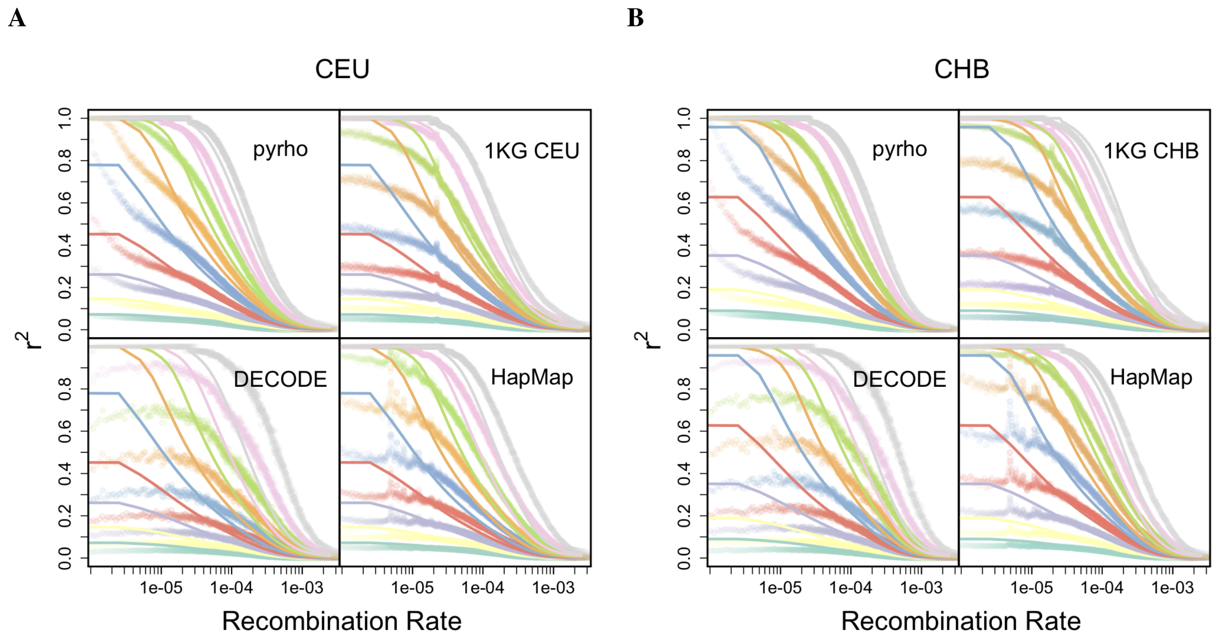


**Figure S3:** Fit of empirical (points) to theoretical (lines) deciles of $r^2$ between pairs of SNPs with MAF greater than 10% as a function of genetic distance between them.

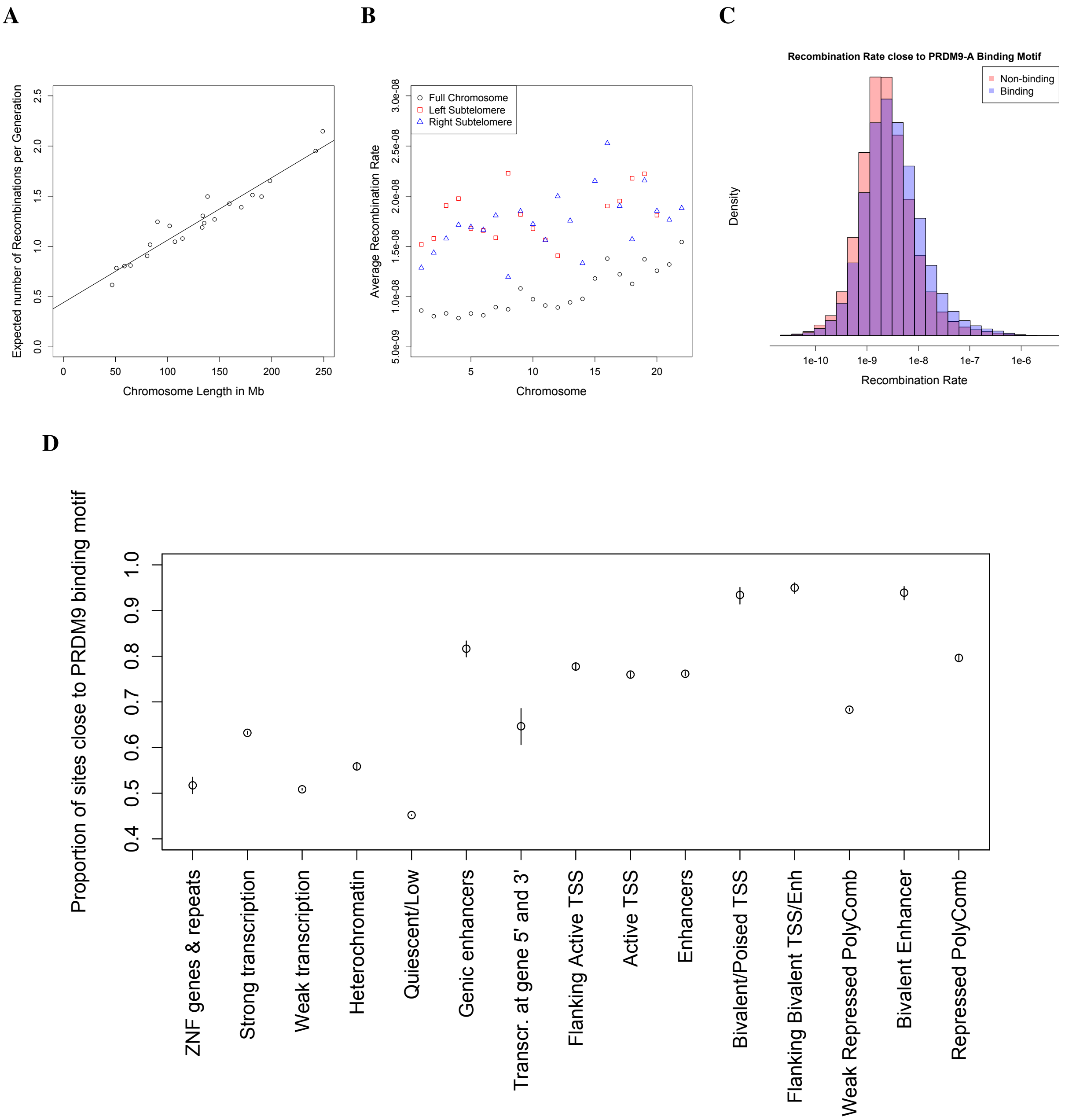


**Figure S4:** (**A**) Expected number of recombinations by chromosome length. The non-zero intercept suggests that there is a minimum number of crossovers required per meiosis, but the positive slope suggests that longer chromosomes can have more than this minimal number. (**B**) Subtelomeres show elevated rates of recombination on all chromosomes. (**C**) Regions within 100bp of a PRDM9-A binding motif have higher recombination rates on average, but the effect explains only a small amount of the variation in recombination rate. (**D**) Sites in some chromatin states are far more likely to be within 100bp of a PRDM9-A binding motif.

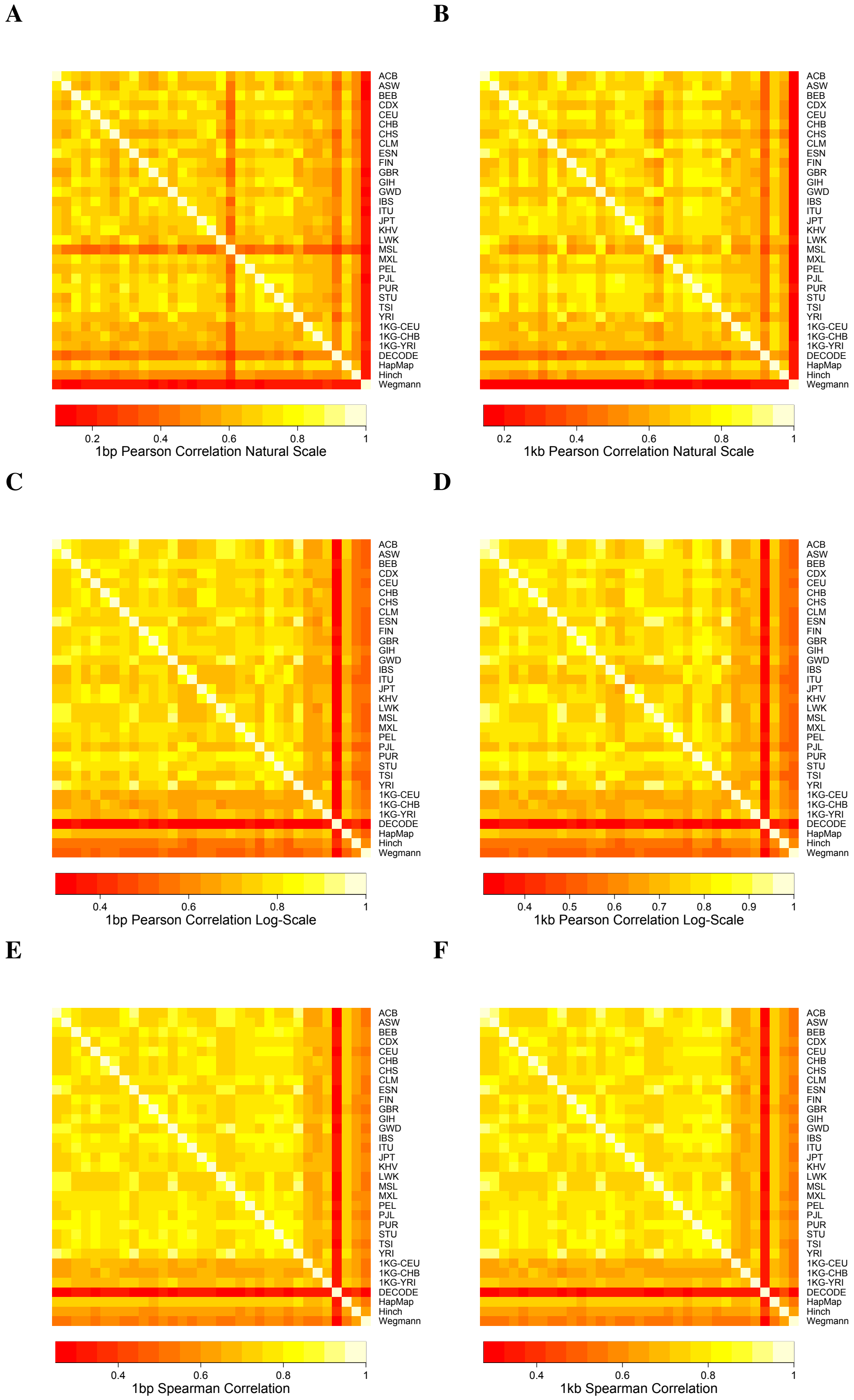


**Figure S5:** Correlation between maps inferred by `pyrho` and maps inferred by previous methods.

**A**

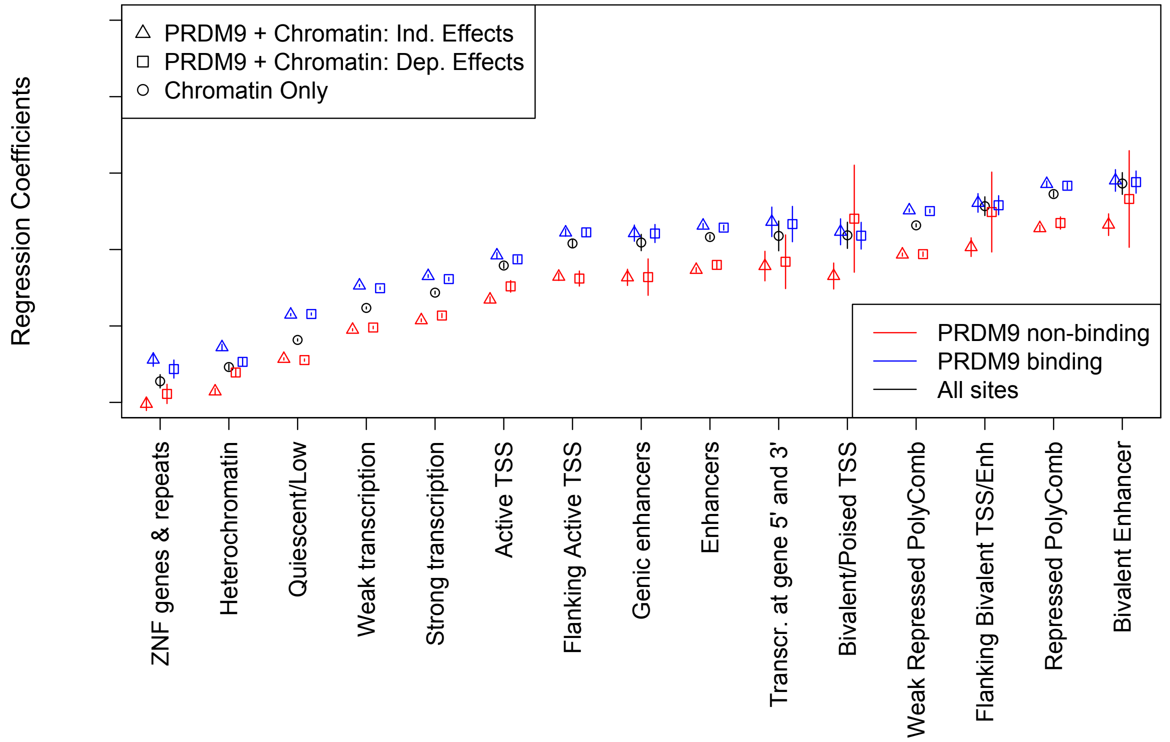


**B**

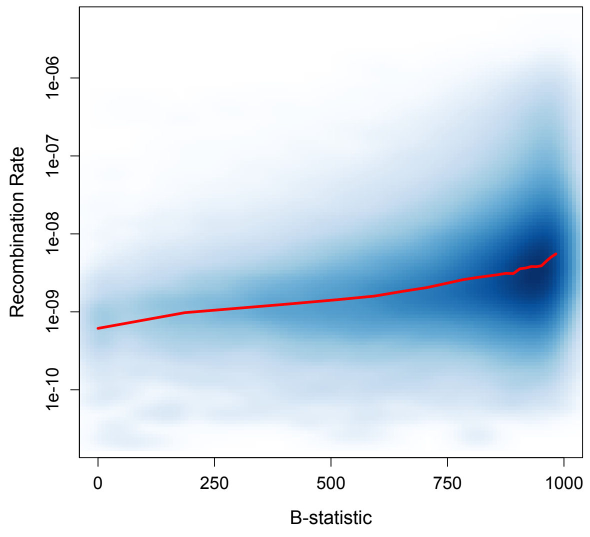


**Figure S6:** (**A**) Inferred regression coefficients after controlling for B-statistics. The results are roughly comparable to Figure 4 in terms of relative ordering and relative effect size. (**B**) Inferred recombination rates as a function of background selection, presented as a smoothed scatter plot with a local average.

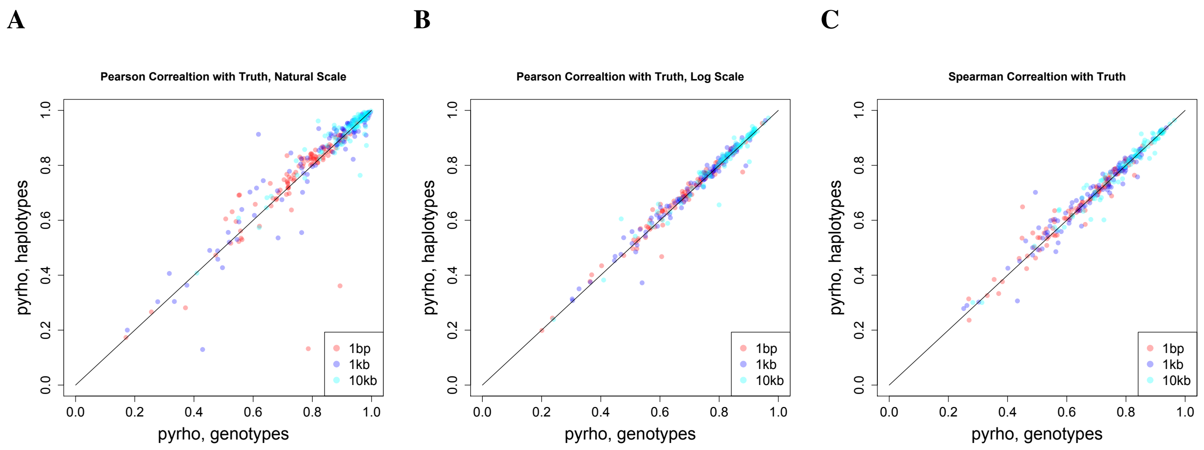


**Figure S7:** Accuracy of simulated data for `pyrho` using phased and unphased data. Accuracy is measured in terms of (**A**) Pearson correlation with the truth in natural scale, (**B**) Pearson correlation with the truth in log scale, and (**C**) Spearman correlation with the truth.

| Corr. | `pyrho` | $\texttt{LDhat}_{\text{demo}}$ | $\texttt{LDhat}_{\text{const}}$ |
|---|---|---|---|
| Pear., 1bp | $\mathbf{0.756 \pm 0.032}$ | $0.536 \pm 0.033$ | $0.471 \pm 0.034$ |
| Pear., 1kb | $\mathbf{0.816 \pm 0.039}$ | $0.778 \pm 0.039$ | $0.746 \pm 0.043$ |
| Pear., 10kb | $0.916 \pm 0.020$ | $\mathbf{0.921 \pm 0.018}$ | $0.864 \pm 0.026$ |
| Pear., log-scale, 1bp | $\mathbf{0.687 \pm 0.013}$ | $0.627 \pm 0.012$ | $0.610 \pm 0.013$ |
| Pear., log-scale, 1kb | $\mathbf{0.713 \pm 0.014}$ | $0.669 \pm 0.124$ | $0.648 \pm 0.013$ |
| Pear., log-scale, 10kb | $\mathbf{0.811 \pm 0.012}$ | $0.808 \pm 0.010$ | $0.791 \pm 0.011$ |
| Spear., 1bp | $\mathbf{0.659 \pm 0.014}$ | $0.605 \pm 0.014$ | $0.597 \pm 0.014$ |
| Spear, 1kb | $\mathbf{0.689 \pm 0.014}$ | $0.653 \pm 0.014$ | $0.640 \pm 0.014$ |
| Spear, 10kb | $0.794 \pm 0.012$ | $\mathbf{0.807 \pm 0.012}$ | $0.799 \pm 0.012$ |

**Table S1:** Correlation (Pearson in natural or log-scale and Spearman) at different spatial resolutions. The mean correlation across 100 simulations ($\pm$ 2 standard errors) is reported, and the best performing method for each measure of accuracy is presented in boldface. We present results for our method, `pyrho`, as well as for `LDhat` using a demography-aware lookup table ($\texttt{LDhat}_{\text{demo}}$) or assuming a constant demography ($\texttt{LDhat}_{\text{const}}$). Overall, the methods that take demography into account outperform $\texttt{LDhat}_{\text{const}}$, and `pyrho` substantially outperforms $\texttt{LDhat}_{\text{demo}}$ at fine-scales, and performs comparably at broader scales.

| Population | $r^{1bp}_{nat. scale}$ | $r^{1kb}_{nat. scale}$ | $r^{1bp}_{log scale}$ | $r^{1kb}_{log scale}$ | $\rho^{1bp}$ | $\rho^{1kb}$ |
|---|---|---|---|---|---|---|
| ACB | 0.814 | 0.818 | 0.997 | 0.997 | 0.997 | 0.997 |
| ASW | 0.877 | 0.859 | 0.997 | 0.997 | 0.997 | 0.997 |
| BEB | 0.855 | 0.867 | 0.986 | 0.987 | 0.987 | 0.987 |
| CDX | 0.958 | 0.967 | 0.988 | 0.988 | 0.989 | 0.989 |
| CEU | 0.828 | 0.826 | 0.991 | 0.991 | 0.992 | 0.992 |
| CHB | 0.863 | 0.843 | 0.984 | 0.985 | 0.986 | 0.986 |
| CHS | 0.916 | 0.925 | 0.985 | 0.985 | 0.986 | 0.986 |
| CLM | 0.840 | 0.825 | 0.991 | 0.990 | 0.992 | 0.991 |
| ESN | 0.742 | 0.748 | 0.998 | 0.998 | 0.998 | 0.998 |
| FIN | 0.986 | 0.987 | 0.990 | 0.990 | 0.990 | 0.990 |
| GBR | 0.980 | 0.982 | 0.990 | 0.990 | 0.990 | 0.990 |
| GIH | 0.916 | 0.913 | 0.989 | 0.990 | 0.990 | 0.990 |
| GWD | 0.492 | 0.499 | 0.994 | 0.994 | 0.994 | 0.995 |
| IBS | 0.920 | 0.904 | 0.988 | 0.988 | 0.990 | 0.990 |
| ITU | 0.787 | 0.799 | 0.985 | 0.986 | 0.986 | 0.986 |
| JPT | 0.904 | 0.900 | 0.987 | 0.987 | 0.988 | 0.988 |
| KHV | 0.800 | 0.807 | 0.989 | 0.989 | 0.989 | 0.989 |
| LWK | 0.945 | 0.944 | 0.996 | 0.996 | 0.996 | 0.997 |
| MSL | 0.693 | 0.585 | 0.996 | 0.997 | 0.997 | 0.997 |
| MXL | 0.984 | 0.942 | 0.991 | 0.990 | 0.991 | 0.991 |
| PEL | 0.961 | 0.885 | 0.986 | 0.985 | 0.987 | 0.986 |
| PJL | 0.928 | 0.914 | 0.992 | 0.984 | 0.990 | 0.991 |
| PUR | 0.836 | 0.837 | 0.994 | 0.994 | 0.995 | 0.995 |
| STU | 0.474 | 0.475 | 0.984 | 0.984 | 0.986 | 0.986 |
| TSI | 0.925 | 0.895 | 0.987 | 0.986 | 0.987 | 0.987 |
| YRI | 0.920 | 0.927 | 0.994 | 0.994 | 0.995 | 0.995 |

**Table S2:** Correlation between maps inferred on hg38 and those inferred on hg19 and lifted over to hg38. Pearson correlation is denoted by $r$ with the subscript denoting whether it is in log-scale or natural scale. Spearman correlation is denoted by $\rho$. The amount of smoothing performed is denoted by the superscript.
